# Identification and Pharmacological Characterization of Multiple Allosteric Binding Sites on the Free Fatty Acid 1 Receptor

**DOI:** 10.1101/2024.12.09.627639

**Authors:** Siddhant Chaturvedi, Krishna Venka, Rahul Singh, Mohandas Kumar

## Abstract

Free fatty acid receptor 1 (FFA1 or GPR40), activated by medium- and long-chain fatty acids, amplifies glucose-stimulated insulin secretion, making it a promising target for type 2 diabetes. Radioligand studies revealed distinct binding sites for partial and full agonists, with full agonists showing positive cooperativity. Functional assays demonstrated positive cooperativity between agonists and varying interactions with the endogenous fatty acid DHA. These findings suggest three allosterically linked binding sites on FFA1, with activation influenced by key arginine residues. Potent ligands with strong cooperativity hold significant therapeutic potential.

## INTRODUCTION

The FFA1 receptor (GPR40) is closely related to other fatty acid receptors, FFA2 and FFA3, sharing 30–50% sequence homology, particularly within transmembrane domains. Free fatty acids, essential for cellular function, also act as signaling molecules.^1^ FFA1 is activated by medium- and long-chain fatty acids, coupling preferentially to Gq to stimulate phospholipase C activity. The receptor is expressed in the brain, monocytes, and pancreatic β-cells, where it plays a role in glucose-dependent insulin secretion. While some studies suggest FFA1 is significant in type 2 diabetes pathogenesis, recent findings challenge this claim.^2,3^ Transgenic expression of GPR40 in β-cells improves glucose tolerance and insulin secretion under a high-fat diet. Synthetic FFA1 ligands, including clinical candidates TAK-875 and AMG 837, have shown promise as therapeutic agents. In this study, novel synthetic agonists displaying full or partial agonism relative to DHA were investigated using radioligand binding assays, revealing three distinct allosterically linked binding sites on FFA1.^4,5^ Key arginine residues, Arg183 and Arg258, were found to be critical for receptor activation and ligand binding. These findings enhance our understanding of FFA1 activation and highlight the therapeutic potential of allosteric agonists for type 2 diabetes.^6^

Functional assays (in vitro, ex vivo, and in vivo) further demonstrated positive functional cooperativity between full and partial agonists.^7-9^ Interestingly, the endogenous fatty acid docosahexaenoic acid (DHA) showed negative or neutral cooperativity with both series of agonists in binding assays but exhibited positive cooperativity in functional assays.^10-12^ Another synthetic agonist was found to act allosterically with both agonist series but appeared competitive with DHA. These findings suggest the presence of three allosterically linked binding sites on FFA1, with each site being selectively targeted by specific agonists.^12-15^

Mutational analyses revealed that activation of FFA1 by these agonists is differentially influenced by mutations in two arginine residues previously identified as critical for FFA1 binding and activation. The high potency of these ligands, coupled with their strong positive functional cooperativity with endogenous fatty acids, demonstrated both in vitro and in vivo, underscores their potential to deliver significant therapeutic benefits.^16-20^

## MATERIALS AND METHODS

The full-length human FFA1 gene was amplified by PCR from human universal cDNA and subcloned into the mammalian expression vector pIRESHyg3 (Clontech). The aequorin DNA was subcloned into the pcDNA3.1 vector (Invitrogen). Chinese hamster ovary (CHO) cells were stably transfected with both FFA1 and aequorin constructs, creating a double-stable cell line maintained in Dulbecco’s modified Eagle’s medium/nutrient mixture F-12 (DMEM/F-12; Cellgro) supplemented with 10% fetal bovine serum (FBS), antibiotics, and hygromycin (600 μg/ml).

A stable FFA1-overexpressing cell line was developed by retroviral infection of A9 cells using a retroviral vector (PLPC) encoding full-length human FFA1. This cell line was cultured in DMEM/F-12 supplemented with 10% FBS, antibiotics, and puromycin (2 μg/ml). Additionally, a cell line stably expressing FFA2 was generated by transfecting CHO cells stably expressing aequorin with the bicistronic expression plasmid pIRES (Clontech) encoding the FFA2 receptor, using Fugene 6 (Roche Diagnostics).

These cell lines were maintained under specific selective conditions to ensure stable expression of the desired constructs.

### Immunohistochemistry

For the localization analysis of FFA1 and GLP-1, paraffin-embedded sections of small intestines from FFA1−/−/β-Gal KI mice underwent heat-induced antigen retrieval in DIVA antigen retrieval solution (Biocare). Sections were blocked using Peroxidaze 1 and Background Sniper (Biocare) and incubated sequentially with rabbit anti-β-galactosidase antibody (Abcam, 1:6000) for 30 minutes and Envision+ rabbit HRP polymer (DAKO). Signal development was performed with DAB+ (DAKO), and selected microscopic fields were photographed.

Subsequently, the same sections were incubated with rabbit anti-GLP-1 antibody (Abcam, 1:2000) for 30 minutes, followed by MACH2 rabbit AP polymer (Biocare). The color reaction was developed using the BCIP/NBT substrate system (DAKO), and identical microscopic fields were photographed after both staining steps. Positive cells were counted to identify those stained for β-gal, GLP-1, or both.

To confirm co-localization, a CRi Nuance multispectral imaging camera (Caliper Life Sciences) was used to analyze the double-stained slides. This technology identifies and localizes chromogens by their spectral signatures, enabling precise detection of overlapping antigens within the same cells. Results were visualized in pseudofluorescence, where β-gal expression appeared green, GLP-1 expression appeared red, and co-localized cells were represented in yellow. These findings demonstrated co-expression of β-gal and GLP-1 in specific cells of the duodenum.

### Test Compounds

Long-chain fatty acids (Sigma–Aldrich) were dissolved in either DMSO or ethanol, while AM-6331 and AM-8182 were dissolved in DMSO to prepare 10 mM stock solutions. Serial dilutions of the test compounds were performed to assess dose-response effects in aequorin and inositol phosphate (IP) accumulation assays. The final volume of test compound solutions added to the assays did not exceed 0.1% for the aequorin assay and 1% for the IP assay, ensuring minimal solvent interference with assay results.

### Aequorin Assay

CHO cells stably expressing both FFA1 and aequorin DNA were seeded into 15 cm dishes and harvested 24 hours later using 2 mL of 1× trypsin-EDTA (0.25% trypsin and 21 mM EDTA in Hank’s Buffered Salt Solution (HBSS); Mediatech Inc.). Cells were pelleted by centrifugation at 600g for 5 minutes and resuspended in HBSS supplemented with 0.01% fatty-acid-free human serum albumin (HSA) and 20 mM HEPES (Cellgro). The suspension was incubated with 1 μg/mL coelenterazine (PJK GmbH, Kleinblittersdorf, Germany) and test compounds at room temperature for 2 hours.

Ligand-induced receptor activation and calcium release were assessed by measuring aequorin luminescence using an EG&G Berthold 96-well luminometer. The luminescence response was recorded over a 20-second interval following the addition of test compounds to the cells. The FFA2 aequorin assay followed the protocol described by Lee et al. (2008).

### Animals and Diets

All animal protocols were approved by the AMGEN Institutional Animal Care and Use Committee. Experimental cohorts were group-housed in humidity- and temperature-controlled rooms with a 12-hour light/dark cycle and provided chow (Teklad 2018S) and water ad libitum. Unless otherwise specified, all mice used in the experiments were male and 2–4 months old.

FFA1-deficient (FFA1−/−) and GPR120-deficient (GPR120−/−) mice were generated by Lexicon Pharmaceuticals (The Woodlands, TX). Breeding pairs for each strain were shipped to Amgen South San Francisco, where colonies were maintained by backcrossing founders to C57BL/6 (wild-type) mice from The Jackson Laboratory (Bar Harbor, ME). Experimental cohorts, including wild-type and knockout littermates, were derived from heterozygous matings and confirmed to be congenic to the C57BL/6J background using microsatellite marker analysis.

GLP-1 receptor knockout (GLP-1R KO) mice were obtained from Dr. Daniel Drucker’s laboratory (University of Toronto, Canada). Microsatellite marker analysis confirmed congenicity ranging from 97% to 100%. Experimental cohorts were produced from homozygous matings, with age-matched C57BL/6J mice serving as controls.

### Data Analysis

Results from in vivo experiments are presented as means ± SEM, with statistical significance determined by either two-way ANOVA or Student’s unpaired t-test, as specified in the figure legends. Data from in vitro pharmacology studies are expressed as means ± SEM of at least three independent experiments. Ligand-induced FFA1 activation is represented as a percentage increase from the basal (unstimulated) level.

Aequorin and inositol phosphate (IP) accumulation assay data were analyzed using the αβ operational model (Leach et al., 2007), with non-linear regression analysis performed in Prism 5.01 (GraphPad Software Inc., San Diego, CA) with modifications to fit the specific experimental parameters. This approach ensured robust interpretation of ligand activity and receptor pharmacology.

## RESULTS

### Activation of the FFA1 Receptor by Synthetic and Endogenous Agonists in Various In Vitro Assays

Novel synthetic agonists for the FFA1 receptor were identified through high-throughput screening followed by detailed structure-activity relationship (SAR) studies. Their specificity, up to concentrations of 30 μM, was verified against a panel of GPCRs, including FFA2, FFA3, GPR119, and glucagon-like peptide 1 receptors. The chemical structures of these synthetic ligands, along with endogenous ligands DHA and α-linolenic acid (LA), are shown in Fig. 1. Among the synthetic agonists tested—AMG 837, AM 8182, and AM 1638—their potencies and efficacies were compared with DHA in various functional assays. AM 1638 and AM 8182 acted as full agonists, while AMG 837 was a partial agonist, achieving approximately 30–50% of the maximum efficacy (Emax) of full agonists across different assays.

**Figure 1.**
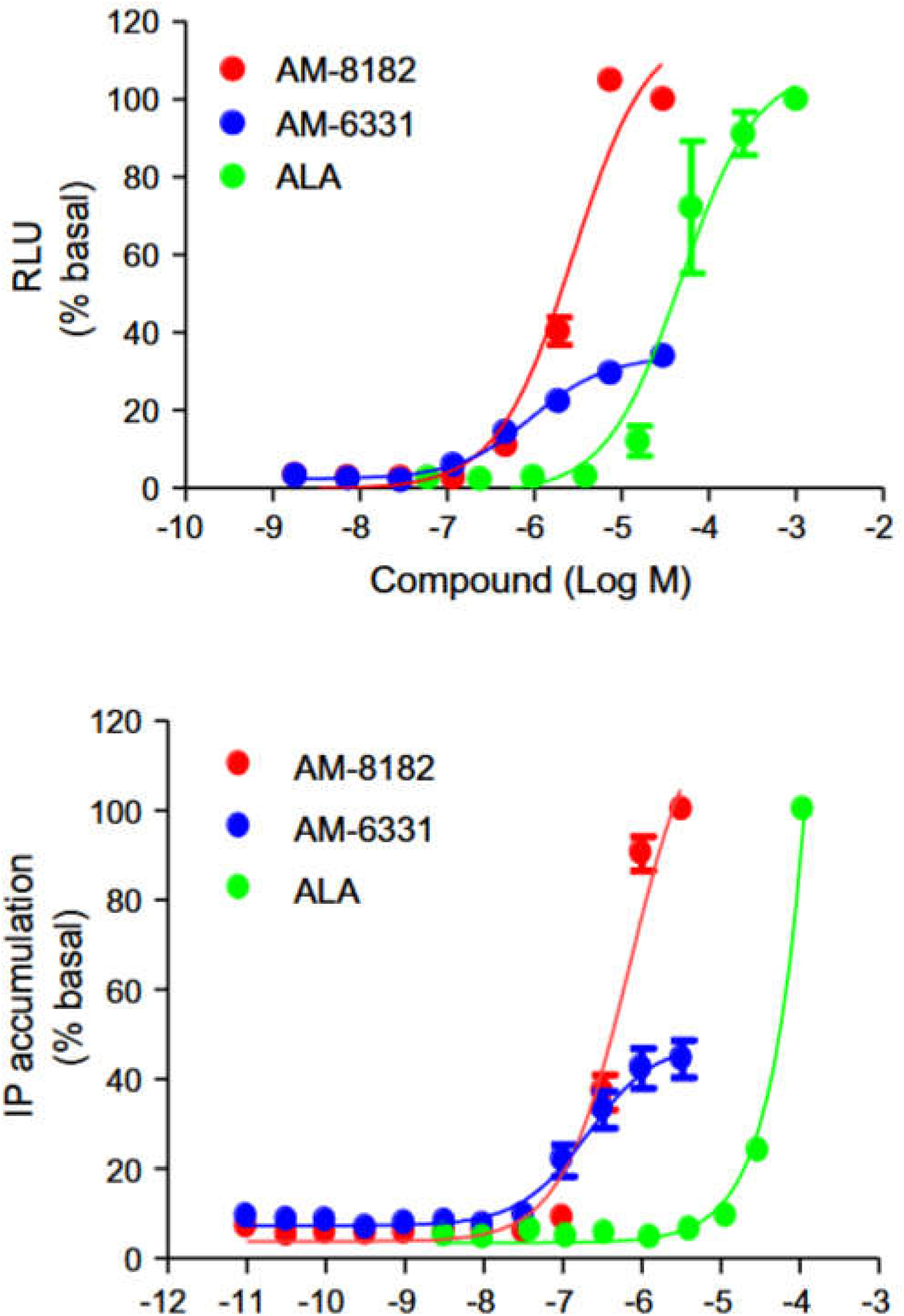
Activity test of the compound related to FFA1

**Figure 2.**
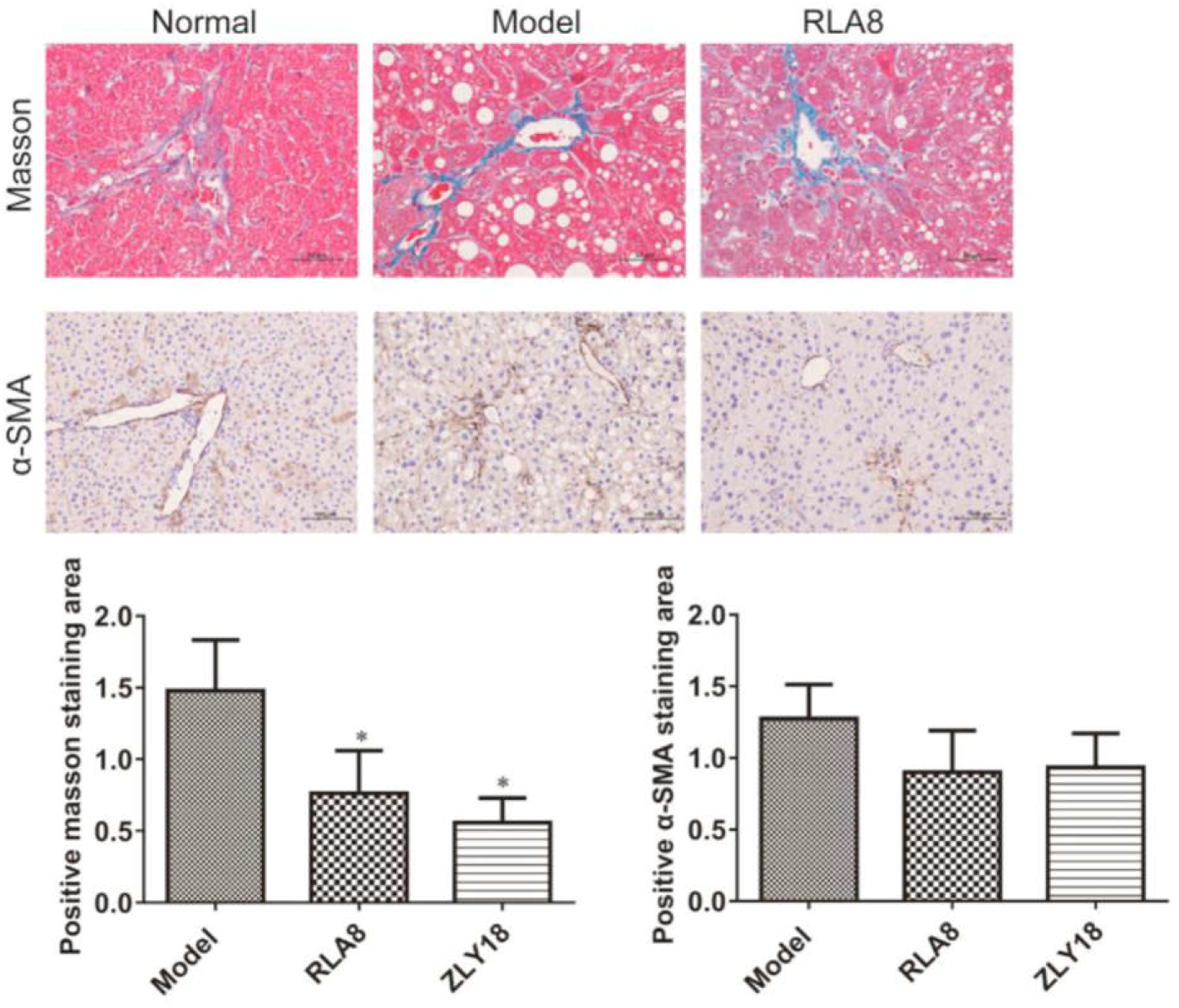
Mice experiments shows the activity in vivo

In aequorin assays conducted in CHO cells expressing low levels of the FFA1 receptor (0.5 pmol/mg protein), AM 1638 and AM 8182 demonstrated full agonist activity, while AMG 837 exhibited partial agonism, with an Emax approximately 30% of the full agonists. Both AMG 837 and AM 1638 were 400–500 times more potent than DHA (EC50 = 40 μM), and AM 8182 was 15-fold more potent. IP accumulation assays in A9 cells with higher receptor expression (6–9 pmol/mg protein) confirmed these findings, showing that AM 8182 and AM 1638 retained full agonist efficacy comparable to DHA, whereas AMG 837 remained a partial agonist, with an Emax only 50% that of DHA and the full agonists. All ligands were 10–40 times more potent in the IP assay than in the aequorin assay, and the synthetic agonists demonstrated 15–1500 times greater potency than DHA across both assays.

These findings illustrate that synthetic FFA1 agonists exhibit significantly higher potency than endogenous ligands, with AM 1638 and AM 8182 showing full agonism and AMG 837 exhibiting partial agonism, making them valuable tools for further exploring FFA1 receptor function and potential therapeutic applications.

### Binding of Radioligands to the FFA1 Receptor

The low-efficacy agonist AMG 837 and the high-efficacy agonist AM 1638 were radiolabeled with tritium, and their binding to A9 membranes expressing the FFA1 receptor was analyzed. Saturation binding curves for both [^3^H]AMG 837 and [^3^H]AM 1638 fit simple binding isotherms (slope factor = 1). The mean log affinities (Kd values) were 8.44 ± 0.05 (3.6 nM, n = 7) for [^3^H]AMG 837 and 7.87 ± 0.03 (13 nM, n = 6) for [^3^H]AM 1638. Both radioligands showed similar Bmax values of 7.9 ± 0.5 pmol/mg protein and 8.4 ± 0.7 pmol/mg protein, respectively, with no significant differences (unpaired t-test).

### Equilibrium Binding Assays of Synthetic Ligands and DHA with the Radiolabeled FFA1 Receptor

The interaction of AMG 837, AM 1638, AM 8182, and DHA with 5 nM [^3^H]AMG 837 was evaluated. As expected, AMG 837 fully inhibited specific [^3^H]AMG 837 binding with a mean pIC50 of 8.13 ± 0.06, consistent with its Kd from saturation experiments. Surprisingly, AM 1638 enhanced [^3^H]AMG 837 binding, indicating positive cooperativity. Data analyzed via the allosteric ternary complex model estimated AM 1638’s log affinity at the unoccupied receptor as 7.58 ± 0.07 and at the [^3^H]AMG 837-occupied receptor as 8.14 ± 0.10, corresponding to a 3.6-fold positive cooperativity.

In contrast, AM 8182 and DHA inhibited [^3^H]AMG 837 binding, but their inhibition was noncompetitive as curves did not extrapolate to zero specific binding at high concentrations. Analysis revealed log affinities of 6.04 ± 0.08 for AM 8182 and 5.33 ± 0.03 for DHA at the unoccupied receptor, with 28 ± 5-fold and 9.8 ± 0.5-fold negative cooperativity, respectively, relative to [^3^H]AMG 837. These results indicate that AM 1638, AM 8182, and DHA bind to distinct sites from AMG 837. Moreover, the binding of [^3^H]AMG 837 and the enhancing effect of AM 1638 were unaffected by guanosine 5′-O-(3-thio)triphosphate (10 μM), confirming their interaction is not dependent on G-protein coupling.

These findings demonstrate the complex allosteric interactions among FFA1 agonists and underscore the potential for allosteric modulation in receptor targeting.

## CONCLUSION

FFA1, a member of the G protein-coupled receptor family A, is mediated by medium- and long-chain fatty acids, leading to the amplification of glucose-stimulated insulin secretion. This highlights FFA1 as a potential therapeutic target for type 2 diabetes. Initially, it was assumed that fatty acids and synthetic FFA1 agonists shared a single binding site. However, radioligand binding studies with members of two distinct chemical series of partial and full agonists have revealed otherwise. The findings show that full agonists bind to a different site than partial agonists, exhibiting positive heterotropic cooperativity.

Functional assays (in vitro, ex vivo, and in vivo) further demonstrated positive functional cooperativity between full and partial agonists. Interestingly, the endogenous fatty acid docosahexaenoic acid (DHA) showed negative or neutral cooperativity with both series of agonists in binding assays but exhibited positive cooperativity in functional assays. Another synthetic agonist was found to act allosterically with both agonist series but appeared competitive with DHA. These findings suggest the presence of three allosterically linked binding sites on FFA1, with each site being selectively targeted by specific agonists.

Mutational analyses revealed that activation of FFA1 by these agonists is differentially influenced by mutations in two arginine residues previously identified as critical for FFA1 binding and activation. The high potency of these ligands, coupled with their strong positive functional cooperativity with endogenous fatty acids, demonstrated both in vitro and in vivo, underscores their potential to deliver significant therapeutic benefits.

This study highlights the distinct and complex binding behaviors of FFA1 receptor agonists, offering insights into their allosteric interactions and therapeutic potential. Radioligand binding assays with [^3^H]AMG 837 and [^3^H]AM 1638 revealed significant differences in binding affinity and efficacy, with AMG 837 acting as a low-efficacy partial agonist and AM 1638 as a high-efficacy full agonist. Notably, AM 1638 exhibited positive cooperativity with AMG 837, whereas AM 8182 and DHA showed noncompetitive inhibition, suggesting distinct and allosterically linked binding sites. These interactions were independent of G-protein coupling, as indicated by the absence of effects from guanosine 5′-O-(3-thio)triphosphate.

The findings confirm that synthetic agonists bind to unique sites on the FFA1 receptor, enabling differential modulation of receptor activity. The positive cooperativity observed with AM 1638 and its potent binding profile underscore its potential as a lead candidate for therapeutic development. Meanwhile, the allosteric mechanisms observed with synthetic and endogenous ligands provide a foundation for the design of targeted therapies leveraging the receptor’s multifaceted binding landscape, particularly in the treatment of type 2 diabetes.

